# Regional fat depot masses are influenced by protein-coding gene variants

**DOI:** 10.1101/526434

**Authors:** Matt J. Neville, Laura BL. Wittemans, Katherine E. Pinnick, Marijana Todorčević, Risto Kaksonen, Kirsi H. Pietiläinen, Jian’an Luan, Robert A. Scott, Nicholas J. Wareham, Claudia Langenberg, Fredrik Karpe

## Abstract

With the identification of a large number of genetic loci associated with human fat distribution and its importance for metabolic health, the question arises as to what the genetic drivers for discrete fat depot expansion might be. To date most studies have focussed on conventional anthropometric measures such as waist-to-hip ratio (WHR) adjusted for body mass index. We searched for genetic loci determining discrete fat depots mass size using an exome-wide approach in 3 large cohorts.

Here we report an exome-wide analysis of non-synonymous genetic variants in 17,212 participants in which regional fat masses were quantified using dual-energy X-ray absorptiometry. The missense variant *CCDC92_S70C_*, previously associated with WHR, is associated specifically with reduced visceral and increased leg fat masses. Allele-specific expression analysis shows that the deleterious minor allele carrying transcript also has a constitutively higher expression. In addition, we identify two variants associated with the transcriptionally distinct fat depot arm fat (*SPATA20_K422R_* and *UQCC1_R51Q_*). *SPATA20_K422R_*, a rare novel locus with a large effect size specific to arm, and *UQCC1_R51Q_*, a common variant exome-wide significant in arm but showing similar trends in other subcutaneous fat depots. In terms of the understanding of human fat distribution, these findings suggest distinct regulation of discrete fat depot expansion.

**Author summary:** Human fat storing tissues are heterogeneous and comprise functionally and structurally distinct regional fat depots, the relative size of which appear to have significant implications for health. Whilst it is known that inter-individual differences in fat distribution have genetic drivers, studies to date have focussed on crude anthropometric approximations of region fat masses rather than precise measures. Here we describe an exome-wide analysis of a large collection of men and women who have undergone body scanning using dual-energy X-ray absorptiometry (DXA) to better define regional fat masses and identify new genetic drivers for human fat distribution. With this approach we identify three gene regions associated with distinct fat depots which can help to explain the variation in fat distribution between people and may lead to a better understanding of the depot specific fat tissue expansion.

## Introduction

Beyond associations with chronic disease and overall obesity, as defined by body-mass index (BMI), it is becoming increasingly apparent that there is an even stronger relationship between body fat distribution and cardio-metabolic disease[1, 2]. For example, Yusuf *et al*.[1] showed that waist-to-hip ratio (WHR) is a stronger predict of myocardial infarction than BMI. To date, the overwhelming majority genome and exome-wide association studies on fat distribution have focussed on waist and hip circumference and WHR[3, 4]. While these measures are easy and cheap to obtain on a large scale, they do not capture all variation in fat distribution. For example, WHR does not capture peripheral fat stored in the upper limbs and the distribution of overall central fat over the subcutaneous and visceral compartments, of which the latter have been suggested to have discordant effects on cardio-metabolic risk[5–8]. Furthermore, circumference-based estimates of fat accumulation do not take into account differences in lean mass and bone structure and mass. Therefore, additional genetic association studies that call upon direct measures of regional fat mass would help unpick mechanisms underlying the expansion of distinct fat depots. Quantification of distinct fat depot masses requires imaging methods with post-image processing to derive delineation of tissues, such as magnetic resonance imaging or dual x-ray absorptiometry (DXA). We have therefore formed a large consortium with DXA-derived regional body fat measurements together with capability to pursue an exome chip discovery project of exonic gene coding variants relating to distinct fat depot size. We hypothesise that by identifying fat depot-specific genetic loci we may gain better insight into the site-specific role of adipose tissue to disease aetiology.

## Results and discussion

We tested the associations of coding genetic variants covered on the Illumina Human Exome Bead chip with regional fat masses measured by DXA (GE Lunar iDXA). Our analyses included up to 17,212 participants of European ancestry from the Oxford Biobank[10], Fenland[11] and EPIC-Norfolk[12] cohorts (Table 1 and Table S1). We fitted within each cohort additive, recessive and dominant models for six DXA-derived adipose tissue regions, i.e., arm fat, leg fat, gynoid fat, total android fat, visceral abdominal fat and subcutaneous abdominal fat (Table S1), using RAREMETALWORKER[13]. The regional fat phenotypes were adjusted for the first 4 principal components, age and total body fat percentage and the residuals were rank-based inverse normally transformed for men and women separately. Meta-analyses of the single variant association statistics were performed in RAREMETAL[14]. Only non-synonymous variants were considered and the cut-off for exome-wide statistical significance was *p*<2E^−7^. Three non-synonymous variants reached exome-wide significance (Fig1, Table 2 and S2 Table): rs11057401, a common missense variant in Coiled-Coil Domain Containing 92 (*CCDC92_S70C_*); rs62621401, a novel low-frequency missense variant in Spermatogenesis Associated 20 (*SPATA20_K422R_*) and rs4911494, a common missense variant in Ubiquinol-Cytochrome C Reductase Complex Assembly Factor 1 (*UQCC1_R51Q_*). An additional 30 non-synonymous variants reached suggestive significance across 38 tests (*p*<10^−6^, S3 Table), including a large haplotype block on chromosome 17 containing 8 missense variants across the *SPPL2C, MAPT, KANSL1* genes and *GDF5_S276A_* in LD with *UQCC1_R5IQ_*.

**Fig 1.**
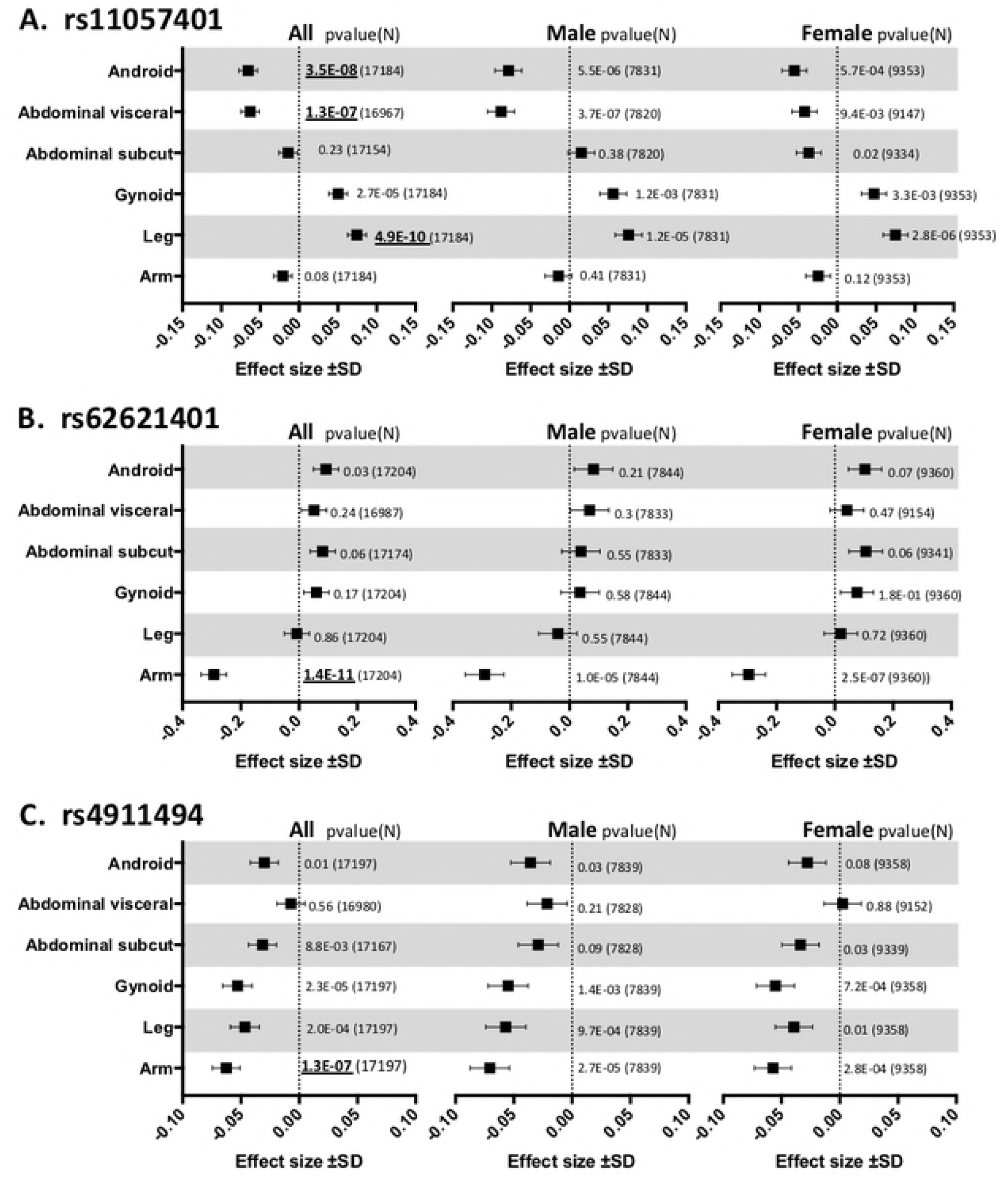
The effect size and direction of effect of Meta-analysis findings. Effect size and direction of effect of the three exome-wide significant missense variants: **A.** rs11057401 in CCDC92 (EAF=0.32), **B.** rs62621401 in SPATA20 (EAF=0.016) and **C.** rs4911494 in UQCC1 (EAF=0.62). Data is presented for the 6 DXA measures under investigation and is presented as the beta value ± SD. The meta-analysis significance level using an additive model for gender combined (All) as well as for gender stratified analysis, together with the N indicated to the right of the data in parentheses. DXA measures are Arm fat mass (Arm), Total android fat mass (Android), Subcutaneous android fat mass (Abdominal subcut), Visceral android fat mass (Abdominal visceral), Gluteal fat mass (Gynoid) and Leg fat mass (Leg). Exome-wide significant data (p<2E^−7^) are in bold and underlined.

**Table 1.**
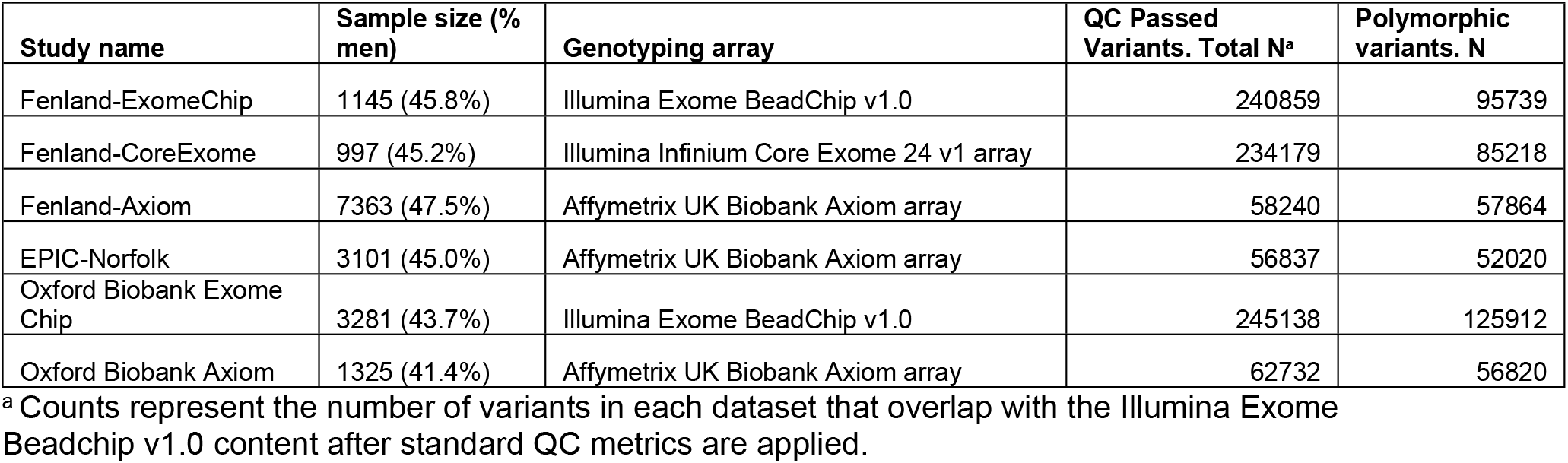
Study Cohorts

| Study name | Sample size (% men) | Genotyping array | QC Passed Variants. Total N<sup>a</sup> | Polymorphic variants. N |
| --- | --- | --- | --- | --- |
| Fenland-ExomeChip | 1145 (45.8%) | Illumina Exome BeadChip v1.0 | 240859 | 95739 |
| Fenland-CoreExome | 997 (45.2%) | Illumina Infinium Core Exome 24 v1 array | 234179 | 85218 |
| Fenland-Axiom | 7363 (47.5%) | Affymetrix UK Biobank Axiom array | 58240 | 57864 |
| EPIC-Norfolk | 3101 (45.0%) | Affymetrix UK Biobank Axiom array | 56837 | 52020 |
| Oxford Biobank Exome Chip | 3281 (43.7%) | Illumina Exome BeadChip v1.0 | 245138 | 125912 |
| Oxford Biobank Axiom | 1325 (41.4%) | Affymetrix UK Biobank Axiom array | 62732 | 56820 |

<sup>a</sup> Counts represent the number of variants in each dataset that overlap with the Illumina Exome Beadchip v1.0 content after standard QC metrics are applied.

The rs11057401 *CCDC92_S70C_* variant (EAF=0.32) is predicted to cause a deleterious amino acid change as assessed by predictSNP[15]. The minor allele of rs11057401 shows significant opposing effects on android, specifically visceral fat mass, and lower body fat, in a sex-combined additive model (total abdominal fat mass= β −0.065, SD 0.0119, *p*=3.5E^−8^; visceral abdominal fat mass= β −0.063, SD 0.0119, *p*=1.3E^−7^; leg fat mass= β 0.075, SD 0.02, *p*=4.9E^−10^) (Table 2). These data extend WHR associations reported by Justice, *et al*.[3] and Lotta *et al*.[11] at this locus. Lotta *et al*.[11] describe the contribution of leg fat-mass; here we demonstrate an additional opposing effect specifically on abdominal visceral fat mass but not for abdominal subcutaneous fat mass, which would correspond to the already observed association with increased waist circumference but with the present analysis showing that the effect is confined to the intra-abdominal fat depot only. Found on chromosome 12q24, *CCDC92* is ubiquitously expressed with highest levels in adipose tissue, brain and testes. It is a nuclear protein interacting with the centriole-ciliary interface[11] and may also be involved in DNA repair[16]. The lead variant tags a large haplotype of at least 99 SNPs (r^2^>0.9) across a number of genes including the putative transcription factor Zinc Finger Protein 664 (*ZNF664*) and Dynein Axonemal Heavy chain 10 (*DNAH10*) (Table 2). There is also strong evidence for multiple eQTL signals across this haplotype which includes three genes, i.e. *CCDC92, DNAH10* and ZNF664[17]. Previous GWAS studies have also associated SNPs in this haplotype with a reduction in insulin resistance[11], improvements in metabolic syndrome[18], reduced WHRadjBMI[3, 4], increased adiponectin levels[19] and with increased plasma HDL-cholesterol and reduced triglyceride concentrations[20–22]. Ablation of *CCDC92* and *DNAH10* in mouse OP9-K cells impairs adipogenesis and reduced lipid accumulation[11]. To further define the likely causative gene or genes in this complex region we undertook a number of gene expression studies in human regional adipocytes and whole adipose tissue. *CCDC92* and *ZNF664* showed very similar expression profiles between abdominal subcutaneous (ASAT), gluteal subcutaneous (GSAT) and arm fat (Fig 2 and 3, Fig S1-3), whilst *DNAH10* expression could not be detected in cDNA from either of the diverse human adipose tissues or cultured primary human preadipocytes making it an unlikely target. Across a panel of 52 paired ASAT and GSAT cDNA samples (Fig S1), qPCR showed small differences in expression of *CCDC92* and *ZNF664* between ASAT and GSAT as well as between lean and obese individuals. In a cultured human primary preadipocyte differentiation time course experiment, both *CCDC92* and *ZNF664* showed a significant upregulation by day 4 of differentiation (Fig 2 A and B) but no difference in expression levels was observed between preadipocytes of ASAT and GSAT origin. Whilst this study focusses on exonic coding variants, recent studies have highlighted the need for caution when dissociating the analysis of such variants from surrounding eQTL signals[23]. To that end we also sought to investigate the reported eQTL signals at this locus, for both *CCDC92* and *ZNF664* (GTEx project[17] and[11] using allele-specific qPCR; a method that allows us to assess expressed allelic imbalance in heterozygous individuals and thus an eQTL. This showed a highly statistically significant increased expression of transcripts found on the minor allele haplotype for both genes (ASAT 5.8%, GSAT 4.9%, Fig 3 A and B). Of functional importance is that this allelic expression imbalance would result in the increased expression of the predicted deleterious serine-70-cysteine amino acid substitution in the *CCDC92* protein. Interestingly, zinc finger proteins such as ZNF664 have been suggested to regulate the expression of near-by genes[18]. The observed co-regulatory expression pattern of *ZNF664* and *CCDC92* could then possibly be due to the eQTL acting on ZNF664 which then upregulates the CCDC92 deleterious variant. Further work needs to be done to investigate this.

**Fig 2.**
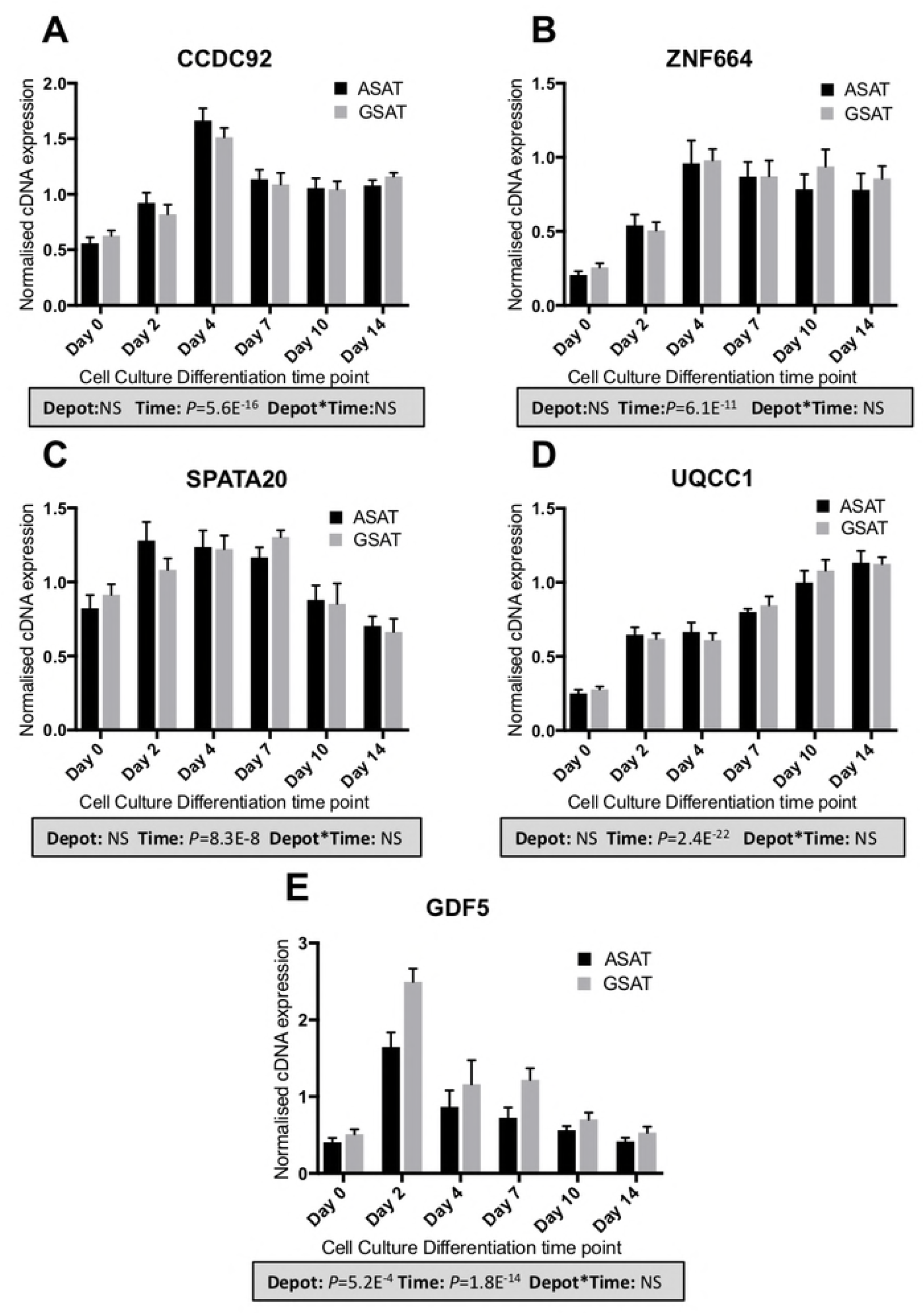
Expression of candidate genes across a human primary adipocyte differentiation time course. cDNA expression of *CCDC92* (**A**), *ZNF664* (**B**), *SPATA20* (**C**), *UQCC1* (**D**) and *GDF5* (**E**) was measured over a 14-day adipogenic differentiation time-course using primary preadipocytes from abdominal subcutaneous (ASAT) and gluteal subcutaneous (GSAT) fat depots[37]. Data are shown as DDCt values (normalized to *PPIA* and *PGK1;* n=6, mean ± SEM). A multivariate general linear model was used to test for statistical significance between depots and time, and to assess depot x time interactions. p-values are presented in the shaded boxes, NS: non-significant.

**Fig 3.**
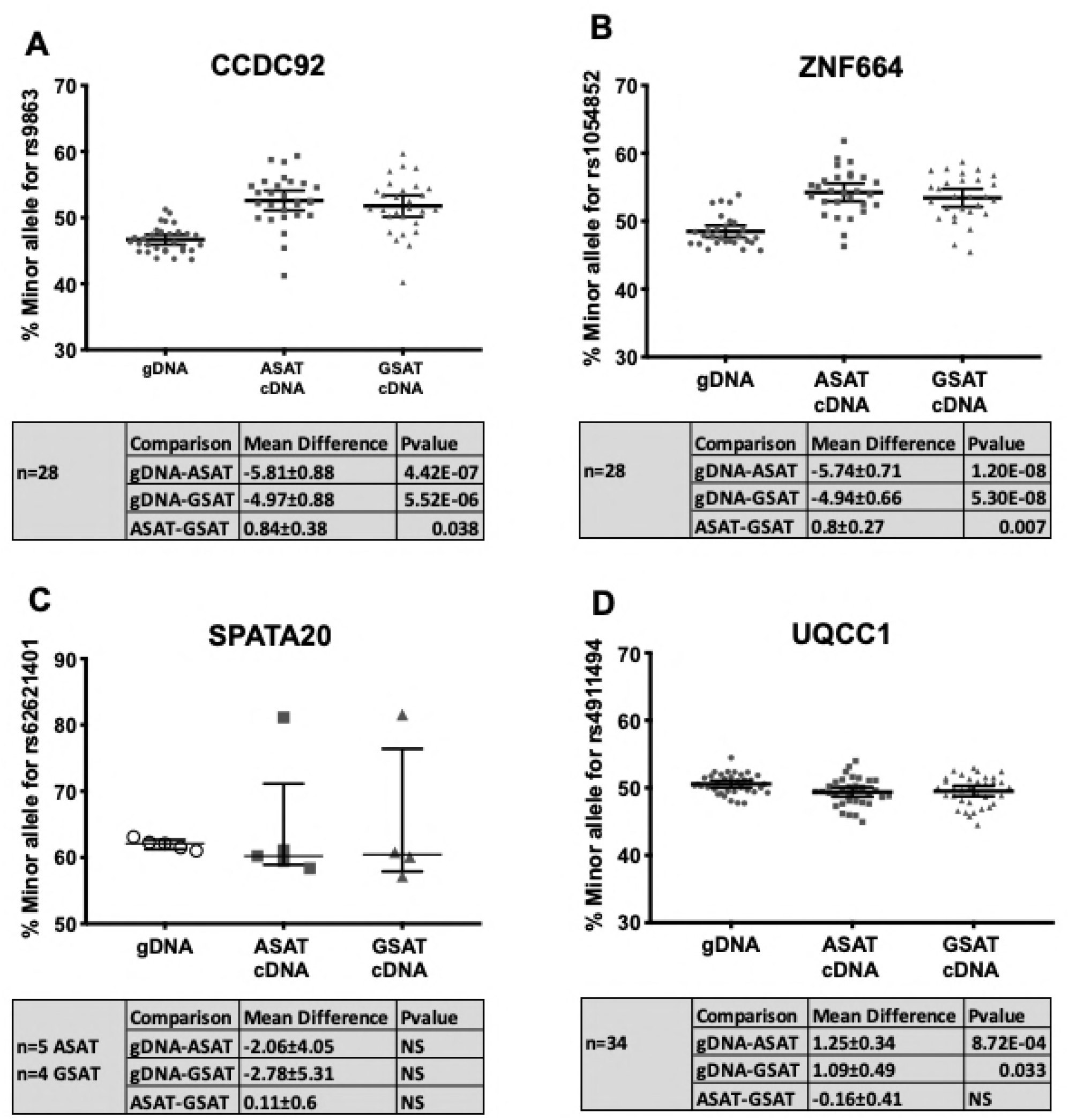
eQTL assessment of exome-wide significant loci by allele-specific qPCR expression. Allelic expression was measured on 4 candidate transcripts in our three exome-wide significant regions using allele-specific qPCR. Data is presented as the % of the minor allele detected compared to the major allele, as described in the methods, with a line indicating the mean and 95%CIs. To assess the rs11057401 eQTL haplotype the proxy SNP rs9863 was assessed for *CCDC92* (A) and the transcribed region proxy SNP rs1054852 for *ZNF664* (B). The index SNP rs62621401 was used to assess the *SPATA20* transcript (C) and the index SNP rs4911494 for *UQCC1*. Paired samples were compared between abdominal subcutaneous (ASAT) and gluteal subcutaneous (GSAT) and genomic DNA (gDNA). For each transcript ABI Taqman genotyping assays were selected that fall within the transcribed sequence. gDNA selected from the same individuals as the cDNAs acts as a paired control with presumed equal allele expression. Deviation from 50% for gDNA, particularly pronounced in *SPATA20* (**C**), represents inherent imbalance in assay technical performance and position of optimal Ct between Vic and Fam fluorescence. By using paired gDNAs to selected cDNAs allelic expression imbalance can be resolved by comparing cDNA to its paired gDNA. Significance was assessed with paired t-test in SPSSv24. Mean differences between comparisons and statistical significance is presented in shaded boxes. NS: Non-significant. The single outlier seen for SPATA20 (C) was replicated in a second cDNA synthesis and both ASAT and GSAT. No phenotype differences were observed for this individual and no obvious genetic differences were observed.

**Table 2.** Primary Exome-Wide significant findings

| rsID | Chr:Position<br>(GRCH37) | Gene | Amino acid<br>Change | Ref<br>Allele | Alt<br>Allele | DXA derived Fat<br>Depot | N | Alternate Allele<br>Frequency | Effect Size | Effect Size SD | Pvalue<sup>a</sup> | Haplotype region<br>(GRCH37)<sup>b</sup> | Number of SNPs in<br>LD with index SNP<sup>c</sup> |
| --- | --- | --- | --- | --- | --- | --- | --- | --- | --- | --- | --- | --- | --- |
| rs11057401 | 12:124427306 | CCDC92 | S70C | T | A | Total Android Fat | 17184 | 0.321 | -0.065 | 0.012 | 3.5E-08 | chr12:124403769-124495203 | 99 |
|  |  |  |  |  |  | Visceral Fat | 16967 | 0.321 | -0.063 | 0.012 | 1.3E-07 |  |  |
|  |  |  |  |  |  | Leg Fat | 17184 | 0.321 | 0.075 | 0.012 | 4.9E-10 |  |  |
| rs62621401 | 17:48628160 | SPATA20 | K422R | A | G | Arm Fat | 17204 | 0.016 | -0.293 | 0.043 | 1.5E-11 | chr17:48623825-48633043 | 7 |
| rs4911494 | 20:33971914 | UQCC1 | R51Q | C | T | Arm Fat | 17197 | 0.616 | -0.063 | 0.012 | 1.3E-07 | chr20:33887955-34194423 | 129 |

<sup>a</sup> Exome-wide significance was set to 2E<sup>-7</sup>
<sup>b</sup> The haplotype region is defined as the furthest 3' and 5' SNPs with a R<sup>2</sup> >0.9 with the index SNP
<sup>c</sup> SNP count is based on 1000 genome SNP data with SNPs in high LD (R<sup>2</sup>>0.9) with the index SNP

The rs62621401 *SPATA20_K422R_* is a rare novel variant (EAF 0.016). This amino acid substitution is not predicted to be damaging and allele-specific qPCR on paired ASAT and GSAT cDNA samples of five Oxford Biobank participants heterozygous for rs62621401 did not reveal any suggestive eQTL (Fig 3C) either. This variant shows a large effect size and is the first locus to be associated with arm fat mass (Arm fat = β −0.29, SD 0.043, p-value=1.5E^−11^, Table 2 and Fig 1B) with an estimated per-allele effect size in the Oxford Biobank (n=4,606) of 125g less arm fat mass (−5.8%, CI −9.3% to −2.3%) in men and 67g (−2.6%, CI −5.3% to 0.4%) in women (approximate fat mass (grams) per allele after adjusting for covariates with % change in parentheses, Table S1). *SPATA20*, linked to spermatogenesis in mice[24], is highly expressed in human testes but also ubiquitously expressed, including in adipose tissue. *SPATA20* is a putative member of the thioredoxin family and other members of this family have been shown to be involved in preadipocyte proliferation[25] and pro-adipogenic *Wnt* signalling[26]. *SPATA20* expression was higher in men than women, although this was only significant for GSAT (p=0.01) (Fig S1). Expression of *SPATA20* during adipocyte differentiation showed an increase between days 0 to 7 of adipogenesis then a drop back to pre-differentiation levels between days 7 and 14 (p=8.3E^−8^, Fig 2C) suggesting a role for this gene in adipocyte development. Surprisingly, despite the arm-specific association, expression of *SPATA20* was similar between arm fat, ASAT and GSAT (Fig S2 A). However, qPCR assessment of a number of developmental genes (Homeobox genes) in arm fat compared to ASAT and GSAT showed significant differences (Fig S2 B) indicating that arm fat is developmentally distinct from the other fat depots assessed. It is therefore possible that SPATA20 is involved in an arm-specific developmental pathway.

The rs4911494 *UQCC1_R51Q_* (EAF=0.62) variant was also associated with a loss of arm fat mass (Table 2 and Fig 1 C) but not predicted as damaging by predictSNP. Whilst exome-wide significance is only observed for arm fat, there is a trend towards less fat in all peripheral and subcutaneous fat depots in both genders (Fig 1 C and Table S1) for the effect allele as described here. Although it should be noted that the minor allele (MAF=0.38) would be associated with an increased fatmass. *UQCC1* is involved with mitochondrial respiratory chain complex III protein expression[27] and is structurally similar to the mouse *Bfzb* controlling mouse brown fat[28]. Previous associations at this locus include height[29], weight[30], WHRadjBMI[3] and osteoarthritis[31]. Another missense variant in the nearby canonical Wnt signalling gene *GDF5* (rs224331) is in LD with rs4911494 (r^2^>0.9) and reaches suggestive significance with arm fat in women (*p*=8.97E^−07^, SD0.027, β 0.13, Table S3). During adipogenesis, expression of *UQCC1* increases (*p*=2.4E^−22^, Fig 2D) but with no difference between ASAT and GSAT in cultured preadipocytes. Allele-specific qPCR showed that the minor allele (rs4911494) was associated with a small but statistically significant decrease in expression of *UQCC1* in ASAT (per allele percentage change in expression = −1.25%, *p*=8.7E^−4^; Fig 3D). It is unclear whether this small change, if confirmed, would be biologically relevant. *GDF5* expression was not detected in adipose tissue cDNA samples. However, during adipogenesis *GDF5* showed transient expression at day 2, highlighting the possibility that *GDF5* is a regulator of early adipocyte differentiation.

## Conclusion

This study represents the largest exome chip meta-analysis on DXA-derived discrete fat depots masses to date. The value of better-defined fat depot regions is illustrated at the *CCDC92* locus. The *CCDC92_S70C_* variant shows a clear signal for visceral fat mass, but none for the adjacent subcutaneous abdominal fat mass and an opposing effect on lower-body fat mass most clearly observed in the whole leg. This is an important distinction from previous waist and WHR associations. Whether these opposing effects are because genetic variation at this locus has direct opposing adipose tissue mass effects in the depots or one depot is simply compensating to the mass change in the other is unclear and will require further investigation. A previous study investigating computerised tomography (CT) scan-derived visceral and subcutaneous fat mass also found associations at the *CCDC92* locus with *ZNF664* (rs1048497) and *DNAH10* (rs1316952)[32] but these SNPs are both in low LD with the index variant in this study (r^2^ 0.34 and 0.18, respectively) and may represent independent signals. No other loci associated with CT-derived visceral fat measures[32] were replicated to suggestive significance. These data and the depth of previous GWAS findings at the *CCDC92-ZNF664* locus highlight this as an important region in regulating adipose tissue distribution. In addition, we report two coding variants associated with arm fat: a novel low-frequency variant in *SPATA20* with a large effect size that seems to only effect arm fat and a common variant in *UQCC1* that additionally seem to have weak effects in other subcutaneous fat depots. Overall, when comparing these data to equivalent exome analysis using anthropometric measures[3] there were few replicated loci. Whilst this lack of replication is likely to be partly due to a lower power compared with significantly larger datasets using conventional anthropometric measures, we have identified a locus, not found for any traditional anthropometric traits, for arm fat and refined the tissue-specific association for another locus (*CCDC92*), highlighting the value of more defined regional fat measures.

Regional fat depots have distinct physiological regulation with an impact on the whole body metabolic homeostasis, with distinct transcriptomes demonstrating functional differences and differences in origin[7]. This study provides genetic evidence for overall, distinct regulation of regional fat depot sizes.

## Materials and methods

### Population cohorts

#### Oxford biobank

The Oxford Biobank (OBB) cohort (http://www.oxfordbiobank.org.uk) consists of an age-stratified random sample of apparently healthy men and women (aged 30 to 50 years) of European ancestry resident in Oxfordshire, UK, as described previously[10]. All participants gave written, informed consent to participate, and studies were approved by the Oxfordshire Research Ethics Committee (08/H0606/107+5). A total of 3,281 individuals from the Oxford Biobank had both measures of fat mass with GE Lunar iDXA[33] and Illumina Human Exome Beadchip genotypes after QC checks. An additional 1,325 individuals contained DXA data and Affymetrix UK Biobank Axiom array genotype data from which overlapping ExomeChip data was extracted for the purposes of this study (Table 1).

#### Fenland

The Fenland study is a population-based cohort study of participants without diabetes born between 1950 and 1975. Participants were recruited from general practice surgeries in Cambridge, Ely and Wisbech (UK) and underwent detailed metabolic phenotyping and genome-wide genotyping.

A total of 1,145 individuals from the Fenland cohort had both measures of fatmass with GE Lunar iDXA[33] and Illumina Human Exome BeadChip genotypes after QC checks (Fenland-ExomeChip, Table 1), a further 997 had Illumina Infinium Core Exome data and 7,363 had Affymetrix UK Biobank Axiom array genotype data from which overlapping ExomeChip data was extracted for the purpose of this study (Table 1).

#### EPIC-Norfolk

EPIC-Norfolk is a prospective cohort study of individuals aged between 40 and 79 years and living in Norfolk county in the UK at the time of recruitment. EPIC-Norfolk is a constituent cohort of the European Prospective Investigation of Cancer (EPIC). A total of 3,101 individuals had Affymetrix UK Biobank Axiom array genotype data from which overlapping ExomeChip data was extracted for the purposes of this study (Table 1).

### DXA-derived depot-specific fat mass measures

For all cohorts’ depot-specific fat mass was quantified using GE Lunar iDXA (GE Healthcare, Bucks, UK). As previously described[33] these give high precision estimates of body composition. The standard setting of the Encore software (version 14.0; GE Healthcare, Bucks, UK) was used to automatically define regions of interest ensuring that boundaries were consistent between cohorts. The descriptives for the DXA measures used are presented in Supplementary Table 1. Visceral fat mass and android subcutaneous fat mass were not measured directly. Visceral fat mass was calculated using an algorithm within the Encore software as described elsewhere[33, 34] and the android subcutaneous fat mass was calculated by subtracting the visceral fatmass from total android fat mass. The DXA scanning was calibrated as per manufacturer’s instructions.

### Exome-wide genotype analysis

#### Datasets

Six data sets from three cohorts, Oxford Biobank[10], Fenland[35] and EPIC-Norfolk[12] (Table 1), equalling a total of 17,212 individuals of European ancestry were compiled for this analysis. The Illumina Exome BeadChip v1.0 genotype content was used as the base content. Where other genotype arrays were used (see Table 1) only the content overlapping with the Illumina ExomeChip were selected. The breakdown of descriptives for each of the 6 datasets can be found in S1 Table. Standard quality control (QC) metrics were employed on each dataset separately and individuals and loci that failed QC removed before association analysis.

#### Single-Variant analysis

All DXA-derived phenotypes were log-transformed, adjusted for age, first 4 principal components (PCs) and percentage total fat mass (calculated as the percentage of total fat mass (grams) to total mass (grams)) and the residuals inverse normal transformed in the R statistical environment. Percentage total fat mass adjusted for age and PC1-4 was also included in the analysis to assess collider bias. Individual datasets were analysed separately in sex-combined and sex-specific analyses using RAREMETALWORKER[13] (http://genome.sph.umich.edu/wiki/RAREMETALWORKER). To account for cryptic relatedness, kinship matrices were first calculated and added into the analysis. Single-variant analysis was performed with, additive, recessive and dominant models.

#### Meta-analysis

Meta-analysis was carried out centrally using RAREMETAL[14]. Variants were excluded of they had a call rate <90%, Hardy-Weinberg equilibrium p-value <1E^−7^ and markers on Y chromosome or mitochondrial genome. Exome-wide significance for the single-variant analysis was set, based on the full ExomeChip content, as *p*<2E^−7^. A suggestive significance was set to *p*<E^−6^.

For this analysis we focussed on non-synonymous variants only, therefore all non-coding variants and synonymous variants were filtered out post meta-analysis. The exome-wide significant findings are presented in Fig 1 and S1 Table; the additional suggestive significant findings are presented in S3 Table.

### Additional informatics

For the three exome-wide significant loci the amino-acid substitutions was assessed for functional significance using the predictSNP online consensus tool[15] (https://loschmidt.chemi.muni.cz/predictsnp1/). This allows for assessment across a number of different tools to generate a consensus assessment. For *CCDC92* the S70C missense variant was assessed; for *UQCC1* the R51Q was assessed and for *SPATA20* three different proteins as products of different splice variants were assessed (K422R, K406R and K362R).

### Adipose tissue gene expression panels

Six genes found within the three index SNP LD boundaries (Table 2) were assessed for expression levels across a collection of human adipose tissue gene expression panels. Applied Biosystems Taqman assay-on-demand qPCR assays were selected for each gene that also avoid the index SNPs presented here, for *CCDC92* (ABI assay, hs01556139), *ZNF664* (ABI assay, hs00921074), *DNAH10* (ABI assay, hs1387352), *SPATA20* (ABI assay, hs00256188), *UQCC1* (ABI assay hs00921074) and *GDF5* (ABI assay, hs00167060).

For tissue panels, subcutaneous adipose tissue biopsies were collected by needle biopsy as previously described[36]. For cell-cultured human primary preadipocytes, of both abdominal subcutaneous fat (ASAT) and gluteofemoral fat (GSAT) origin, a differentiation time course (n=6) was performed as described in Todorčević, *et al*.[37]. All biopsies and cells were homogenized in Tri-reagent (cat. no. T9424, Sigma-Aldrich, UK) and RNA was extracted with a standard Tri-reagent protocol. A total of 500ng RNA was used for cDNA synthesis following standard protocols and random hexamer primers using the cDNA Reverse Transcription Kit (Life Technologies, UK). Real-time PCR reactions were performed on a 1/40 cDNA dilution using Taqman Assays-on-Demand (Applied Biosystems) and Kapa Probe Fast Mastermix (Kapa Biosystems) in triplicate in a 6μl final volume and run on an Applied Biosystems 7900HT machine. Expression was assessed within each panel using a relative qPCR approach[38] and normalised using the previously assessed stably expressed endogenous control genes[36]. For the Lean/Obese Oxford Biobank panel (S1 Fig) the geometric mean of *PPIA, PGK1, PSMB6* and *IPO8* were used. *IPO8* was not used in a paired arm, ASAT and GSAT panel (S2 Fig) as it was not stably expressed between arm and the other depots. *PPIA* and *PGK1* were used as endogenous controls for primary cell culture experiments.

Neither *DNAH10* or *GDF5* could be detected above background in whole tissue cDNA panels. *GDF5* was however detected in a 14-day in vitro adipocyte differentiation time course.

Data for a panel of 52 paired ASAT and GSAT biopsy samples was used to assess expression between sexes, between ASAT and GSAT fat depots, and between lean and obese individuals. Descriptives for this panel are presented in S1 Fig. As both *SPATA20* rs62621401 and *UQCC1* rs4911494 were associated with arm fat mass their expression, along with *CCDC92* and *ZNF664* was assessed in a paired arm, ASAT and GSAT cDNA panel. As there is no published data on arm fat transcriptomics the additional *HOX* gene transcripts *HOXA5, HOXB8, HOXC8, HOXC9* and *HOXC11* were assessed as these are known to be differentially expressed between ASAT, GSAT and visceral fat (These data are presented in S2 Fig).

The setup of a human primary adipocyte differentiation time course is described elsewhere[37]. Relative qPCR was run as above on the adipocyte panel for *CCDC92, ZNF664, SPATA20, UQCC1* and *GDF5*. Data is presented in Fig 2.

### Allele-specific qPCR

Both the *CCDC92* and the *UQCC1* loci are associated with multiple eQTL signals. Whilst we only consider non-synonymous variants in this analysis this does not discount that the coding locus is also under the influence of an eQTL. To assess the available data from resources such as the GTEx portal and to assess any eQTL effect between ASAT and GSAT fat depots we used the combined resources available within the Oxford Biobank.

Allele specific pPCR was run essentially as described in Fogarty *et al*.[39]. Taqman genotyping assays (Applied Biosystems) were selected to fall within the transcripts under investigation. For *CCDC92* the index SNP assay performed poorly so the Proxy SNP rs9863 (ABI assay, C_206415_30) was selected. To assess the nearby gene *ZNF664* a SNP in high LD with the CCDC92 index SNP that fell within the *ZNF664* transcript, rs1054852 (ABI assay, C_1169873_10), was selected. For *SPATA20* the index SNP was used (rs62621401, ABI assay C_25983779_10) as was for *UQCC1* (rs4911494, ABI assay, C_25472999_10). As was previously stated neither *DNAH10*, nor *GDF5* could be detected in whole adipose tissue panels. Therefore, allele specific qPCR could not be assessed for these two genes.

From a panel of 200 paired ASAT and GSAT cDNA samples available from the Oxford Biobank, heterozygous individuals were selected. For *CCDC92* and *ZNF664* 28 paired ASAT and GSAT samples were selected, for *UQCC1* there were 34 and for *SPATA20* there were 5. Genomic DNA (gDNA) for these individuals were also retrieved and diluted to 1.5ng/μl. The gDNA is used as the control comparison to the cDNA samples as there is an equal quantity of both alleles in heterozygous gDNA samples. By comparing the ratio of the Ct values from each allele (the ratio of the genotype assay Vic or Fam fluorophore signals) between cDNA and gDNA any allelic expression differences observed in the cDNA samples can be resolved. This is particularly relevant as technical variation exists with each genotyping assay; particularly pronounced in *SPATA20* (Fig 3C).

Data are presented as the percentage of the minor allele Ct value compared to the major allele Ct. This is calculated by first generating a standard curve and regression statistic for each assay. A standard curve is generated from genomic DNA for individuals homozygous for the major allele (BB) and minor allele (bb). Genomic DNAs are diluted to 1.5ng/μl then BB and bb homozygotes are combined to ratios 80:20, 60:40, 50:50,40:60,80:20.

Following qPCR analysis using the dual-labelled TaqMan Genotyping assays the ration of the B to b Ct values are calculated (Ct B minus Ct b) then plotted against the percentage of the minor allele in the dilution series. The linear regression statistic from this standard curve is then used to calculate the percentage minor allele expression of the unknown heterozygous individuals. The standard curves are presented in S3A-D Fig and allele-specific qPCR data for heterozygous individuals are presented in Fig 3.

For *CCDC92, ZNF664* and *UQCC1* there were sufficient cDNAs in the 200 panel, however for *SPATA20* there were only 5 individuals. Therefore, to improve the accuracy of the *SPATA20* analysis, each sample was run in triplicate 4x across the assay plate and the average of all 4 sets of triplicates calculated. A single outlier in the SPATA20 data was followed up in a second cDNA synthesis and persisted in both ASAT and GSAT samples. No phenotype differences were observed for this individual and no obvious genetic differences were found.

### Statistical analysis

Statistical significance was assessed for each experiment in SPSS v24. For estimates of per allele grams fat mass change, log phenotype data was analysed in a general linear model and adjusted for age, PC1-4 and total %fat mass then estimated marginal means were calculated (S1 Table).

## Acknowledgements

We thank the volunteers from the Oxford Biobank (www.oxfordbiobank.org.uk) for their participation. The volunteer recruitment and recall process was funded by the National Institute for Health Research (NIHR) Oxford Biomedical Research Centre (BRC). The views expressed are those of the author(s) and not necessarily those of the NHS, the NIHR or the Department of Health recalling process of the volunteers.

## Supporting information

**S1 Fig. mRNA expression of candidate genes across a lean/obese adipose tissue gene expression panel.** mRNA expression of the genes *CCDC92, ZNF664, UQCC1* and *SPATA20* across a panel of paired abdominal subcutaneous fat (ASAT) and gluteofemoral fat (GSAT) cDNA samples from the Oxford Biobank. The panel consisted of 25 male and 29 female healthy individuals selected for either high or low BMI (Lean male, n=13, age 44.5±0.9 yrs, BMI 22.7±0.3 kg/m^2^, fasting blood glucose 5.2±0.1 mmol/l; Obese males, n=12, age 43.4±1.2 yrs, BMI 34.9±5.2 kg/m^2^, fasting blood glucose 5.6±0.1 mmol/l; Lean females, n=15, age 44±1.0 yrs, BMI 21.2±0.2 kg/m^2^, fasting blood glucose 4.8±0.1 mmol/l; Obese Females, n=14, age 44±1.0 yrs, BMI 33.6±0.6 kg/m^2^, fasting blood glucose 5.2±0.1 mmol/l – data expressed as mean ±SEM). Data are shown as the mean ± SEM DDCt values (normalized to the geometric mean of the endogenous control genes *PPIA, PGK1, IPO8* and *PSMB6)* as described previously[36, 38]. A multivariate general linear model was used to test for statistical significance between gender, fat depots and obesity and to assess interactions. P-values are presented in the shaded box, NS: non-significant.

There were small but significant differences in expression of *CCDC92, ZNF664* and *UQCC1* between fat depots in lean individuals but this difference was lost and expression was significantly reduced, in obese individuals. This is in keeping with a general quiescent state observed in transcripts associated with adipocyte metabolic activity in obesity.

**S2 Fig. mRNA expression of Candidate genes and Homeobox genes across a panel of 22 paired arm, abdominal subcutaneous adipose tissue (ASAT) and gluteofemoral adipose tissue (GSAT).** mRNA expression of the candidate genes **A:** *CCDC92, ZNF664, UQCC1* and *SPATA20* and a selection of developmental *HOX* genes **B:** *HOXA5, HOXB8, HOXC8, HOXC9* and *HOXC11* were determined by real-time qPCR. Data are shown as the mean ±SEM DDCt values (normalized to *PPIA, PGK1* and *PSMB6;* n=22). A univariate general linear model was used to test for statistical significance between depots. P-values for the HOX genes in **B** are presented in the shaded box.

**S3 Fig. Allele specific qPCR standard curves and *CCDC92-ZNF664* regression analysis.** The standard curve and regression statistic used to calculate the percentage minor allele expression with allele-specific qPCR is shown above for *CCDC92* (A), *ZNF664* (B), *SPATA20* (C) and *UQCC1* (D). To quantify any allelic expression imbalance for the four genes a standard curve was generated from genomic DNA for individuals homozygous for the Major allele (BB) and Minor allele (bb). Genomic DNAs are diluted to 1.5ng/μl then BB and bb homozygotes were combined to ratios 80:20, 60:40, 50:50,40:60,80:20 to generate a standard curve. Following qPCR analysis using dual labelled TaqMan Genotyping assays the ratio of the B to b allele Ct values are calculated (Ct B minus Ct b) then plotted against the percentage of the minor allele in the dilution series. The linear regression statistic from this (A, B, C and D above) is then used to calculate the percentage minor allele expression of our unknown individuals. For *CCDC92* (A), *ZNF664* (B) and *UQCC1* (D) three different pairs of homozygote individuals were used to generate each standard curve and a Mean ± SEM plotted for each dilution (A, B and D). For *SPATA20* only one genomic DNA homozygote minor allele individual was available so an error bar cannot be displayed.

As discussed in the main text there was an observed co-regulatory pattern of expression between *CCDC92* and *ZNF664* across different cDNA panels. To assess any correlation between these two genes within the samples, the allele-specific qPCR paired data points were plotted and regression statistic calculated (Graphs E and F). For both ASAT (E) and GSAT (F) there was a significant correlation, further supporting the co-regulatory pattern of expression.

**S1 Table. Population cohort descriptives**.

**S2 Table. Exome-wide significant loci.** Detailed data on the three exome-wide significant loci described. DXA parameters are included for all measures and meta-analysis statistics for the additive model. DXA measures are arm fatmass (Arm), Total android fat mass (Android), Subcutaneous android fat mass (Subcut), Visceral android fat mass (Visceral), Gluteal fat mass (Gluteal) and Leg fat mass (Leg). Effect size data for suggestive exome-wide significance (p<=10^−6^) is shown in bold. Exome-wide significant data (p<2E^−7^) are in bold and underlined.

a. The impact of missense variants were assessed using the PREDICTsnp online consensus tool[15] (https://loschmidt.chemi.muni.cz/predictsnp1/).
b. Approximate fat mass (grams) changes per allele is shown where test reaches suggestive significance and were calculated as marginal means after adjusting for age, PCs1-4 and %fatmass as covariates in a general linear model, implemented in SPSS v24

**S3 Table. Exome-wide loci showing suggestive level of statistical significance.**

Additional non-synonymous loci where statistical tests did not reach exome-wide significance but did reach a suggestive significance cut off of p<=10-6 are included above.

a. Where it reaches suggestive significance the model is shown as Additive (add), Recessive(rec) or Dominant (Dom)
b. The impact of missense variants were assessed using the predictSNP online consensus tool[15] (https://loschmidt.chemi.muni.cz/predictsnp1/).
c. The cluster of 8 Missense SNPs found at the SPPL2C-MAPT-KANSL1 locus on chromosome 17 are part of a single haplotype that extends across ~400kb in this region containing >2300 SNPs (r^2^>0.9), rather than independent signals.

